# Temperature dependence of allelopathy duality and its influence on boreal forest succession-A case analysis of *Picea schrenkiana*

**DOI:** 10.1101/516823

**Authors:** Xiao Ruan, Li Yang, Min-fen Yu, Zhao-hui Li, Ying-xian Zhao, Cun-de Pan, De-an Jiang, Qiang Wang

## Abstract

Global warming in conjunction with various biotic or abiotic interferences has been jeopardizing the ecosystem of boreal forests. By integrating field inspection with experimental simulation, this work comprehensively investigated the allelopathic effects of a key allelochemical 3,4-dihydroxyacetophenone (DHAP) in the exudates of *P. schrenkiana* needles on its seed and seedling growth, endogenous hormone metabolism and antioxidant enzyme activity, identified the existence of DHAP allelopathy duality at a certain temperature with an inflection concentration point (e.g. about 0.25 mM at dark/light temperature of 4/12 °C) as the boundary between promotional and inhibitory effect, and verified that the inflection point of DHAP concentration would inevitably shift to a lower level as temperature increased. Consequently, this paper gives a scientific explanation into the intrinsic mechanism of *P. schrenkiana* degradation due to allelopathy, but also presents a new approach to explore the relationship between forest evolution and global warming.

**Highlight:** A quantitative description on the duality of 3, 4-dihydroxyacetophenone (DHAP) as a promoter or an inhibitor to affect the seed germination, seedling growth and root development of *P. schrenkiana*, as well as the antioxidant enzyme activities and hormone contents.

The new findings of DHAP inflection concentration as boundary to divide the promotional and inhibitory effect of allelopathy which would decrease as environment temperatures rise.

An explanation into the intrinsic mechanism of *P. schrenkiana* degradation due to allelopathy, and a new approach to explore the relationship between forest evolution and global warming.

## Introduction

Allelopathy can be generally defined as a direct or indirect, promotional or inhibitory effect of a plant including microorganism on other plants or own through the release of chemicals in environment (Rice, 1984). There have been a lot of research reports about alleochemical production in woody species ranging from Eucalyptus sp. forest in Australia to boreal conifer forest, tropical forest, temperate forest and sub-desert zone communities (Mallik, 2008). Pellissier and Souto (1999) presented a detailed compilation of more than a hundred tree species with allelopathic activity in boreal forest. It has been recognized that allelochemicals could be released from plants into the environment through several ways such as volatilization, root exudation, decomposition and leaching to interfere the growth of adjacent plants (Subrahmaniam *et al*., 2018). Some allelochemicals could increase cell membrane permeability (Bais *et al*., 2003; Chai *et al*., 2013), inhibit cell division and elongation, damage cell sumicroscopic structure (Teerarak *et al*., 2012; Cheng *et al*., 2016), disturb plant photosynthesis and respiration (Yu *et al*., 2005), affect synthesis of plant endogenous hormones and proteins (Zeng *et al*., 2001; Hu *et al*., 2015), and so on. In consequence, allelopathy would potentially influence the growth and development of plants, succession of plant communities (Cummings *et al*., 2012; Bonanomi *et al*., 2018) and invasion of exotic plant into forest ecosystems (Meiners *et al*., 2012). For the purpose of simulation analysis, Blanco (2007) developed a forest ecological-level model FORECAST in incorporating several aspects of allelopathy to demonstrate its potential consequences. Both field investigation and model simulation indicated that the early stages of plant growth were more fragile and sensitive to the changes in variety and quantity of allelochemicals than the adult stages, as such created a major bottleneck to plant propagation.

Depending on its concentration, an allelochemical can act as either a promoter or an inhibiter to plant growth. Such bidirectional behavior may be called as the duality of allelopathic effect. Our previous investigation on the autotoxic effects of *Picea schrenkiana*, the most representative species in boreal forest, has revealed that 3, 4-dihydroxy-acetophenone (DHAP) leached from *P. schrenkiana* needles could display a similar dual effect on the growth of its seedling (Ruan *et al*., 2011). Specifically, DHAP could act as a promoter at lower concentrations (0.1 mM) but an inhibitor at higher concentration (0.5 mM), and the concentration inflection point turning DHAP action direction would appear about 0.25 mM at 4/12 °C. This inflection point would shift to a lower level as environmental temperature increased, which would generate some crucial effects on the early regeneration process of *P. schrenkiana* in boreal forest (Ruan *et al*., 2016). In spite that the real existence of plant autotoxic duality has been gradually recognized, its temperature dependence and ecological significance to forest evolution still need to be fully understood.

Over the decades, climatic change in cooperation with other environmental stressors has been significantly influencing plant ecological behavior and successive propagation (Reich *et al*., 2016; Vazquez *et al*., 2017). The uncertainty of allelopathy induced by climate warming and its effect on boreal forest regeneration require further study. Focusing on the verification of allelopathy duality and its dependence on temperature, this work comprehensively investigates the allelopathy of DHAP to the regeneration of *P. schrenkiana* in boreal forest. In brief, a series of experiments including seed germination, seedling growth, root cell viability, antioxidant enzyme activities and plant endogenous hormones were conducted to reveal the variation of DHAP allelopathy to *P. schrenkiana* regeneration as environment temperature changed.

## Material and Methods

### Geography, climate and ecosystem of the forest

Boreal forests cross over Eurasian continent and account for about 30 % of the global forest area as displayed in Fig.1A. As one of the largest mountain ranges in central Asia, Tianshan Mountain occupies 800,000 km^2^ between 69°-95° E and 39°-46° N, and stretches close to 2,500 km from southwest to northeast in one of the most arid mid-latitude zones on Earth. The forest on Tianshan Mountain range is located in a limited zone with altitudes between 2,700 m (the thermal tree line) and 1,500 m, and the brown soil is covered by a thick humus layer of litterfall over the years. The climate there is rather wet and warm, annually giving the mean temperature, precipitation, evaporation and relative humidity to be 2 □, 400-600 mm, 980-1,150 mm and 65 % respectively, while the aridity index and the frost-free period are 1.4 and 89 days respectively. This multilayered forest is inhabited by a variety of plants including various trees, shrubs, ferns, grasses, and moss. *P. schrenkiana* mingled with *Larix sibiricain* in the eastern region constitutes the major type of forest on Tianshan Mountain. The dominant species of understory shrubs are *Juniperus pseudosabina* and *Juniperus Sabina*, and the main types of understory bryophytes consist of *Dicranum scoparium* and *Hepnum revolutumare*, while the herbs mainly include *Stellaria songorica* and *Cortusa brother*. In addition, the real-time monitoring of soil temperature with a total of sixty-nine sampling points from five field sites on the northern slops of the Tianshan Mountains showed that during natural seasons for *P. schrenkiana* regeneration from 2003 to 2012, the criterion day and night temperatures were 12 °C and 4 °C for seed germination, and 14 °C and 6 °C for seedling growth, respectively (Ruan *et al*., 2016).

**Fig. 1.**
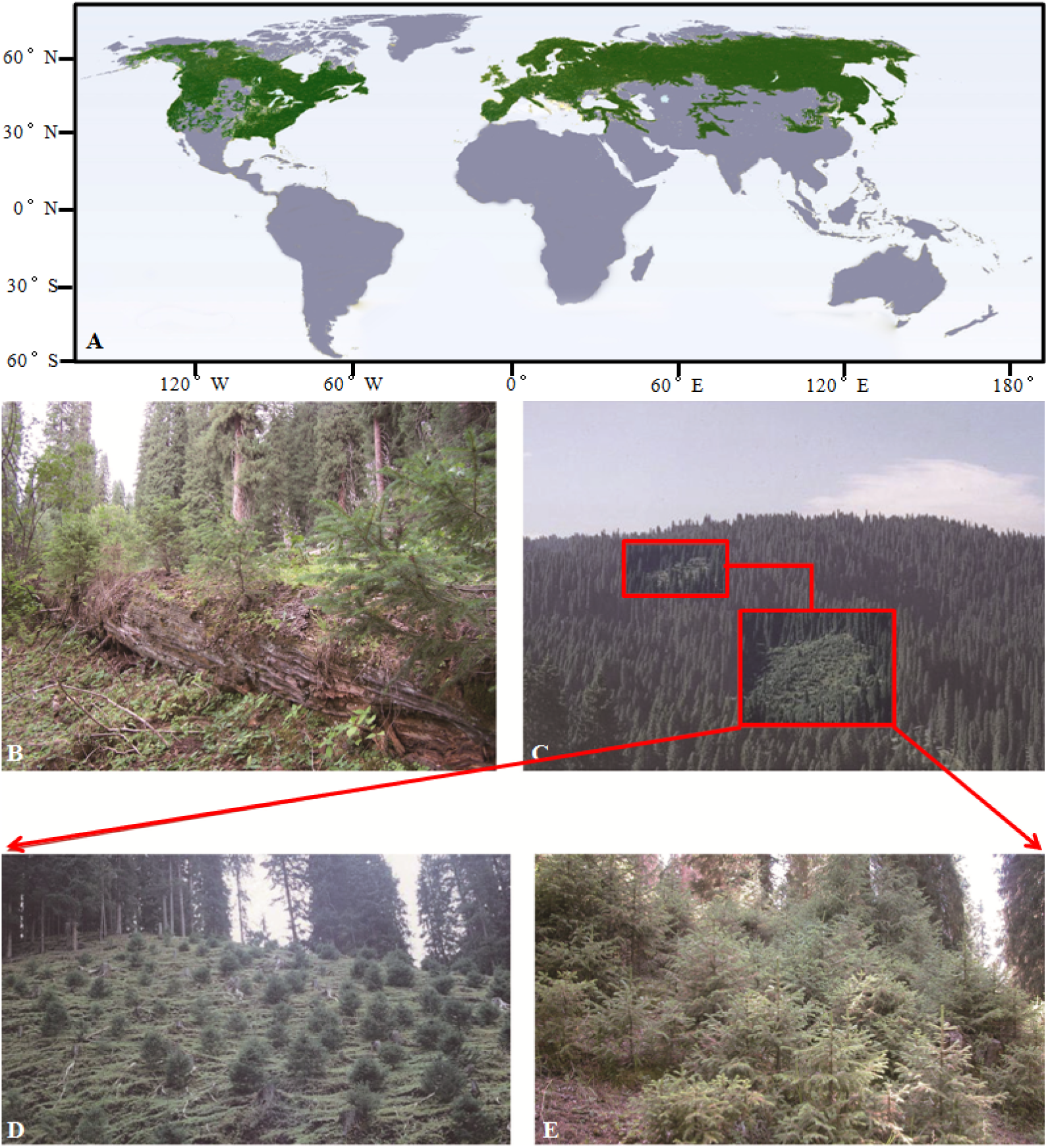
Distribution of boreal forest and regeneration of *P. schrenkiana*. A: Distribution of global boreal forest; B: Regeneration of *P. schrenkiana* in fallen woods (50 years); C: Regeneration of *P. schrenkiana* over sporadic fire spots; D: Regeneration of *P. schrenkiana* after in 5-year after the wildfire event; E: Regeneration of *P. schrenkiana* after in 10-year after the wildfire event

Documentary records indicated that the distribution region and population scale of *P. schrenkiana* were very sensitive to climate change during prehistoric and historic periods. As known for long time, wildfire has played some important role in sustaining natural regeneration and evolution of forest ecosystem. It has been naturally observed that the sustainable circulation of *P. schrenkiana* ecosystem relied on wildfires to some extent (Zhang and Zhang 1963). As shown in Fig.1B, for example, the seedlings and saplings of *P. schrenkiana* only appeared on the burned down woods which experienced the third, fourth and fifth grades of decay respectively, suggesting that the decomposition of the fallen woods could be likely to circumvent or attenuate some negative effects on the seed germination and seedling growth of *P. schrenkiana*. Fig.1C displayed the regeneration of *P. schrenkiana* over two sporadic fire spots, Fig.1D and Fig.1E illustrated the regeneration status of *P. schrenkiana* in five and ten years after the wildfire event occurred respectively. Hereby, a wildfire burning might be appropriate or even beneficial to some forest ecosystems, probably because it could not only damage and even destroy plant morphological landscape, but also attenuate and even eliminate negative or toxic effects of some substances on ecological balance and evolution of forests. Previous investigation showed that water extract of *P. schrenkiana* needles exhibited autotoxic effects on seed germination and seedling growth (Ruan *et al*., 2011; Yang *et al*., 2017).

### Collection of *P. schrenkiana* needles and cones

The current-year needles and cones of *P. schrenkiana* were collected from those parent trees located at the XAU forest education center at 2,198 m altitude, 43°22′58″ north latitude and 86°49′33″ east longitude on September 12-15, 2007-2017. All the selected *P. schrenkiana* plants were 30-35 m tall, 80-100 years old, healthy and infection free. After being collected, the cones were dried in paper bags at room temperature for 7 days and then threshed by hand to get seeds. According to the analyses, the chemical composition of needles and the vigor of seeds showed no difference for the selected plants from various areas (Li *et al*., 2009).

### Extraction and isolation of the active components

A certain amount of dry *P. schrenkiana* needles were ground and extracted with distilled water (20 mL per gram) at room temperature for 48 h. Subsequently, the water solution was extracted again with diethyl ether, ethyl acetate and *n*-butanol in turn. The obtained extracts were concentrated and then fractionated with a silica gel column chromatography to isolate active components. It was previously found that the fractionation of the concentrated diethyl ether extract finally gave a yellow crystal of 3, 4-dihydroxy-acetophenone (DHAP) with the strongest auto-toxicity, and the detailed information of experimental operation and identification has been previously described elsewhere (Ruan *et al*., 2011).

### Bioassay procedure

The stock solution of DHAP at 100 mM was prepared by dissolving pure DHAP in distilled water, and then diluted into the concentrations of 0.5, 0.25 and 0.1mM as treatment solutions and water as control for bioassays. Similarly, the water extraction solution of *P. schrenkiana* needles at 1.0 mg mL^−1^ was prepared by dissolving the water extracts in distilled water, and then diluted to the concentrations of 0.1, 0.05, 0.01 mg mL^−1^ for bioassays. The biological measurements of seed germination and seedling growth were conducted according to the procedure of ISTA (International Seed Testing Association, 1993).

### Measurement of the seed germination

100 seeds of the surface-sterilized *P. schrenkiana* were first put into culture dishes (12×12 cm) lined with two layers of Whatman No 3 filter paper, and each dish was added with 10 mL treatment solution or control water. The seeds were incubated in an artificial intelligence simulation incubator under a 16/8 h (day/night) photo period with photon flux density of 40 μmol m^−2^s^−1^ at a day/night temperature of 8/0 °C, 10/2 °C, 12/4 °C, 14/6 °C and 16/8 °C, respectively. Treatments were conducted in a completely random manner and with five replicates for each. Once the radicle emerged after incubation, the seeds were considered to have germinated. The rate and vigor of germination were calculated after 15 days and on the tenth day, respectively.

### Measurement of the seedling growth

Replicated by five times, one hundred of the successfully germinated seeds were placed in Petri dishes and 10 mL treatment solution was added into each dish, and then the seedlings were incubated in an artificial intelligence simulation incubator under a 16/8 h (day/night) photo period with photon flux density of 40 μmol m^−2^s^−1^ at a day/night temperature of 10/2 °C, 12/4 °C, 14/6 °C, 16/8 °C, and 18/10°C, respectively. After incubation, five seedlings were randomly sampled from each Petri dish, and the length of their shoots and roots was measured with a vernier caliper (GB/T 1214.2-1996, Measuring Instrument LTD, Shanghai). The weight of fresh seedlings was also recorded (Mettler Toledo instrument LTD, Switch). The measurements were taken on the third day after incubation, and continued once every 3 day for a total of 30 days. After their radicle length and fresh weight measured, the seedlings were used to determine the activities of antioxidant enzymes and the levels of plant endogenous hormones.

### Measurement of the root cell viability of seedling

The viability of *P. schrenkiana* root cell was determined by the method of double staining with fluorescein diacetate (FDA) and propidium iodide (PI) (Pan *et al*., 2001). Root tissues (0.1-1 cm length from the tip) were excised from the intact seedlings with or without DHAP treatment, and then the staining process was performed and photographed using a fluorescence microscope (Nikon E600 with a B-2A filter, excitation 450-490 nm, emission at 520 nm, Nikon Corp., Tokyo, Japan) according to the reported methods (Yang *et al*., 2017).

### Assay of plant endogenous hormone contents

Sample of *P. schrenkiana* seedlings (0.1 g) was ground in liquid nitrogen, dissolved with 80% cold methanol (containing 1 mM BHT), and then temporarily incubated at 4□ in the dark for 12 h. After centrifugation at 10,000 r·min^−1^ for 20 min at 4□, the supernatants were prewashed with 80 % methanol, dried under N_2_ and then dissolved in 2 mL methanol for analysis. The contents of plant endogenous hormones were determined by a Agilent 1290 UPLC (Ultra-high Performance Liquid Chromatography) system with a C18 reversed-phase column (2.1×150 mm, Agilent, Santa Clara, CA, USA) in accordance with the method described previously (Yang *et al*., 2017). Zeatin (ZT), Gibberellin (GA_3_), Indoleacetic acid (IAA) and Abscisic acid (ABA) and DHAP as reference materials were assayed by UPLC, and retention time of each compound was measured, as marked in the following bracket: ZT (1.52 min), DHAP (2.95 min), GA_3_ (3.65 min), IAA (5.32 min), ABA (7.35 min), respectively (Fig. 2). The concentrations of plant endogenous hormones (μg·g^−1^ fresh weight) were automatically calculated from peak area by software using authentic standard runs with the sample. All the calibration curves showed excellent linearity (R^2^ >0.999) in a wide concentration range.

**Fig. 2.**
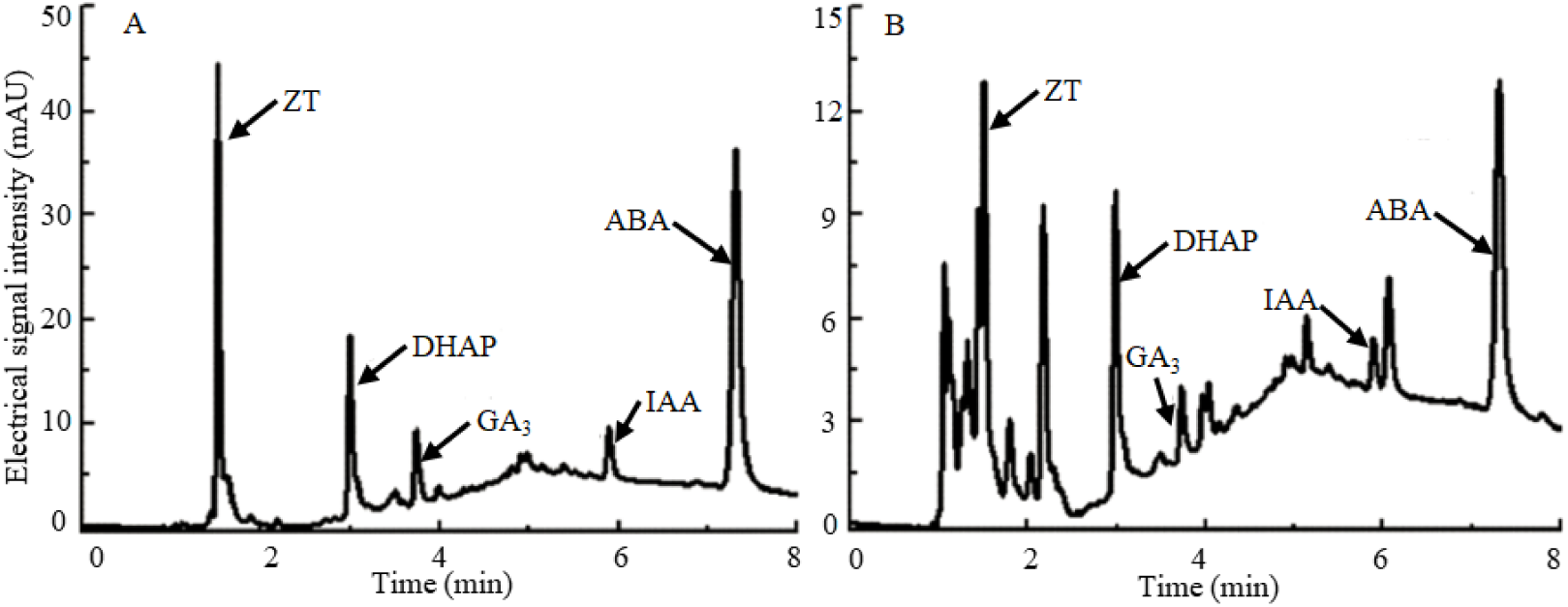
Chromatogram of four phytohormones and DHAP by UPLC. A: Standard chromatogram; B: Sample chromatogram.

### Assay of antioxidant enzyme activities

The antioxidant enzyme activities of *P. schrenkiana* seedling were analyzed by the standard methods and the procedures described previously (Yang *et al*., 2017). In brief, Speroxide dismutase (SOD) activity was determined using the nitrobluetetrazolium (NBT) method, and one unit of the activity was defined as the amount to cause 50 % inhibition of NBT reduction (Giannopolitis and Ries 1977); Peroxidase (POD) activity was detected in accordance with the guaiacol method, and one unit of the activity was defined as the amount of 0.01 increase in the absorbance at 470 nm per min (Kochba *et al*., 1977); Catalase (CAT) activity was determined according to the rate of H_2_O_2_ decomposition as measured by the decrease of absorbance at 240 nm, and one unit of the activity was calculated as the amount of 0.01 decrease in absorbance at 240 nm per min (Zhang *et al*., 1990); Glutathione reductase (GR) activity was assayed by following GSSG-dependent oxidation of NADPH, and one unit of the activity was expressed as 1 μM NADPH oxidized per min (Lee *et al*., 2001).

### Statistical analyses

All results were presented as the mean ± standard error of five replications. All data were statistically analyzed using SPSS software (IBM, New York, USA). For statistical analyses, relationships were considered to be significant when *p*<0.05. If the results of One-way ANOVA showed the significant differences at the 0.05 significance level, LSD (Least Significance Difference) was adopted for multiple comparisons among the different treatments.

## Results

The duality of allelopathy to boreal forest ecosystem depends on both allelochemical concentration and environmental temperature, which can be displayed by investigating the effects of both the allelopathic mixture (water extract) and single key allelochemical (DHAP) in *P. schrenkiana* needles on phenotypic, morphologic and physiological characteristics of *P. schrenkiana* plant at various temperatures.

### Dynamic variability of auto-allelopathic effect on the seed germination

The auto-allelopathic effects of both the water extract of *P. schrenkiana* needles and single DHAP on the germination of *P. schrenkiana* seeds were investigated at three different concentrations and in five dark/light temperature cycles of 0/8, 2/10, 4/12, 6/14 and 8/16 °C. The experimental results (*p*<0.05) are illustrated by a set of diagrams in Fig. 3, in which the vertical axis represents the inhibition ratio as a percentage of the net change value divided by the intrinsic value, so that a positive, negative or zero value indicates an inhibition, promotion or no effect respectively. For either the extract or DHAP, low concentration enhanced the rate and vigor of seed germination but high concentration reduced the rate and vigor at any temperature cycles, while the intermediate concentration could alter the effect from promoting to inhibiting the seed germination as the temperature increased. At a given temperature, therefore, there always was an inflection point of concentration (threshold) for dividing the promotional and inhibitory effect on the seed germination, and increasing the temperature could shift the inflection point to a lower level. For the effect of DHAP on the rate and vigor of seed germination as an example, the inflection concentration point shifted from higher than 0.25 mM to much lower than this level with increasing the dark/light temperature from 0/8 to 4/12 to 8/16 °C (*p*<0.05), as seen in Fig 3C and 3D.

**Fig. 3.**
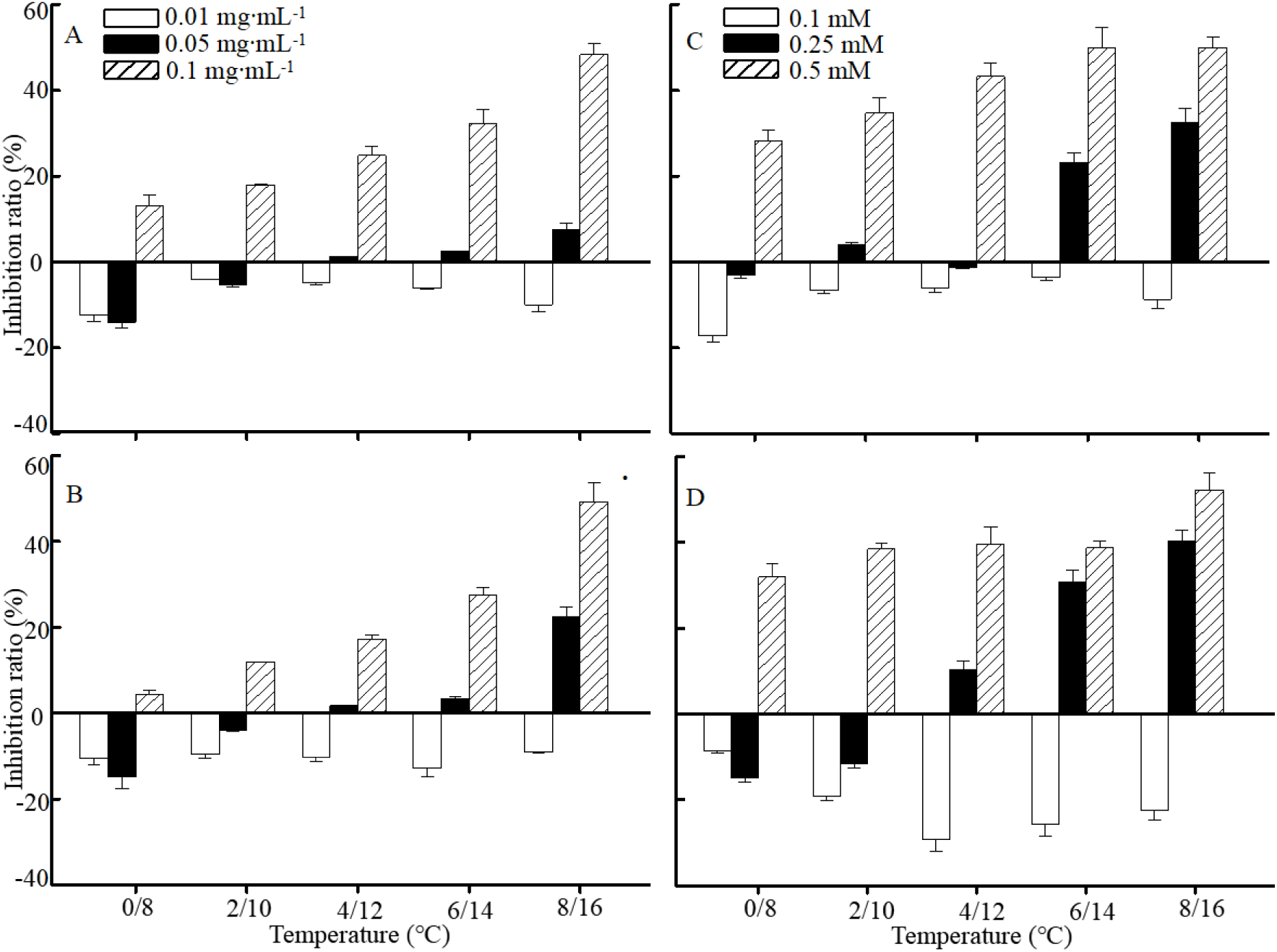
The effect of water extract and DHAP treatment on seed germination of *P. schrenkiana* in different temperature ranges. A-B: Water extract on germination rate and germination vigor; C-D: DHAP on germination rate and germination vigor

### Dynamic variability of auto-allelopathic effect on the seedling growth

Fig. 4 illustrated the experimental results on three different concentrations of the water extract of needles and single DHAP affecting the growth of *P. schrenkiana* seedlings at five dark/light temperature cycles, indicated by the changes of plumule length, radical length and fresh weight (*p*<0.05). Depending on concentration and temperature, the auto-allelopathic effect in each pair of donor (the water extract or single DHAP) and target (plumule length or radical length or fresh weight) could display the duality of promotion or inhibition. Similar to the above seed germination, at any given temperature cycle the low concentration of donor invariably enhanced the growth of *P. schrenkiana* seedlings but the high concentration inevitably reduced the growth, while the intermediate concentration could alter the inhibition ratio from a negative value (promotion effect) through zero line (no effect) to a positive value (inhibition effect) with increasing temperature. For the effect of DHAP on the growth of seedling as an example, the concentration of 0.25 mM could increase the plumule and radical lengths of the seedlings at 2/10, 4/12 and 6/14 °C (*p*<0.05), slightly lengthen the plumule but largely shorten the radicle at 8/16 °C (*p*<0.05), and largely cut down both the lengths at 10/18 °C (*p*<0.05), as showed in Fig 4D and 4E. In brief, there always exists an inflection concentration point of the donor to divide its promotional and inhibitory effect on the growth of *P. schrenkiana* seedlings at a given temperature, and all such temperature-dependent inflection points can be connected into a boundary line that drifts downward with the increase of temperature.

**Fig. 4.**
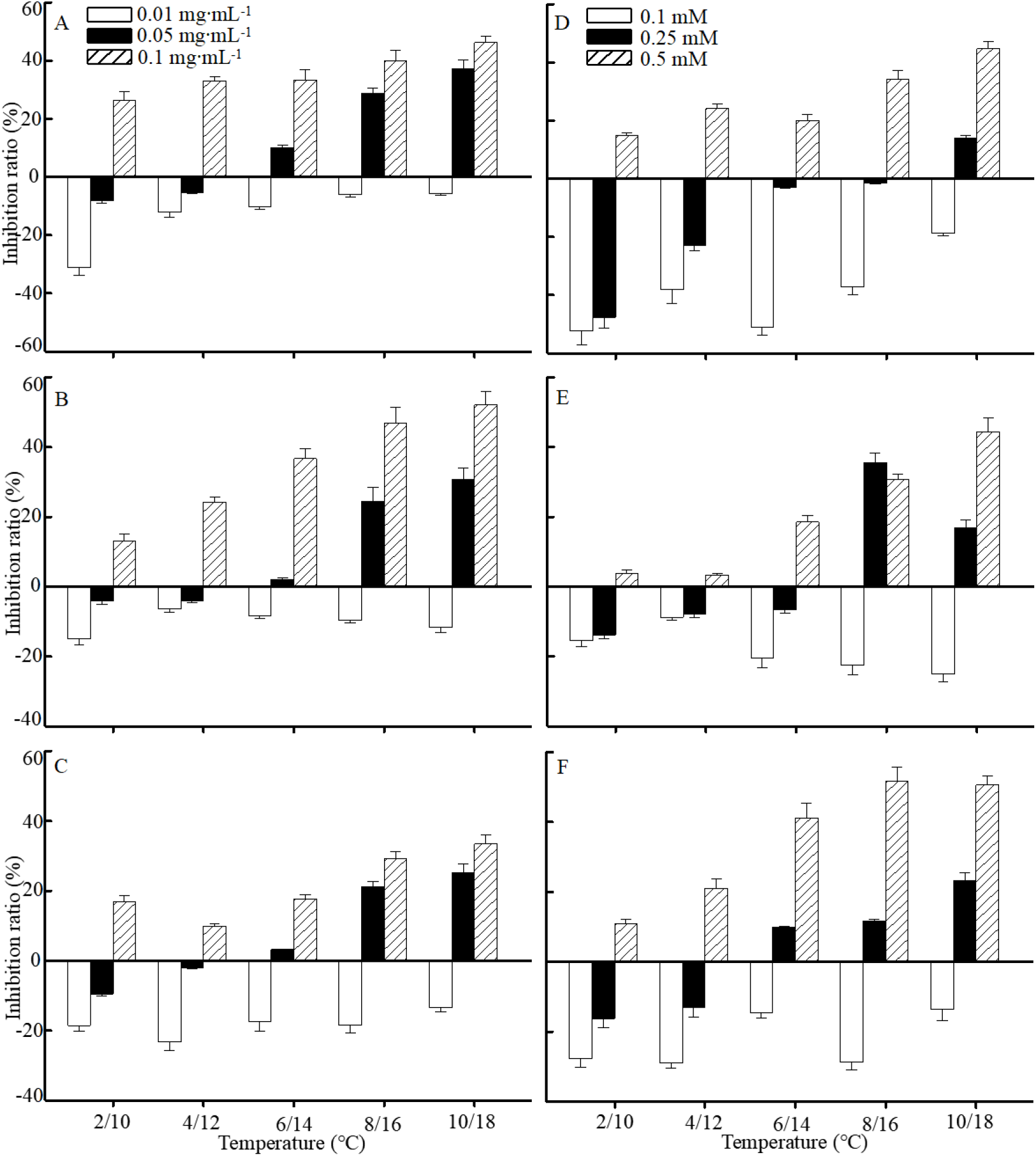
The effect of water extract and DHAP treatment on seedlings growth of *P. schrenkiana* in different temperature ranges. A-C: Water extract on plumule length, radicle length and fresh weight; D-F: DHAP on plumule length, radicle length and fresh weight

### Dynamic variability of auto-allelopathic effect on the viability of root tips

The auto-allelopathic effects of DHAP at three concentrations and five dark/light temperature cycles on the viability of root tips of *P. schrenkiana* seedlings were illustrated by fluorescence in Fig. 5, where fresh green and yellow red indicate alive and dying roots, respectively. As compared with the case in the absence of allelochemical (CK), 0.1 mM DHAP could significantly enhance the vitality of the seedling roots at the dark/light temperature cycles 4/12 and 6/14°C, and 0.25 mM DHAP almost give no effect on the vitality of roots at the temperature below 6/14 °C but a serious damage to the vitality at the temperature over 8/16 °C, while 0.5 mM DHAP cause a loss of roots vitality even at low temperature of 2/10 °C until the complete death of roots at high temperature of 10/18 °C. In general, the relatively low concentration of DHAP at a relatively low temperature could enhance the viability of roots, but the higher concentration of DHAP at a relatively high temperature might kill the roots completely. For the influence of DHAP on the viability of root tips of *P. schrenkiana* seedlings, therefore, there also appears the duality depending on temperature and concentration, although the inflection points are not easy to determine quantitatively and the boundary line is difficult to draw clearly.

**Fig. 5.**
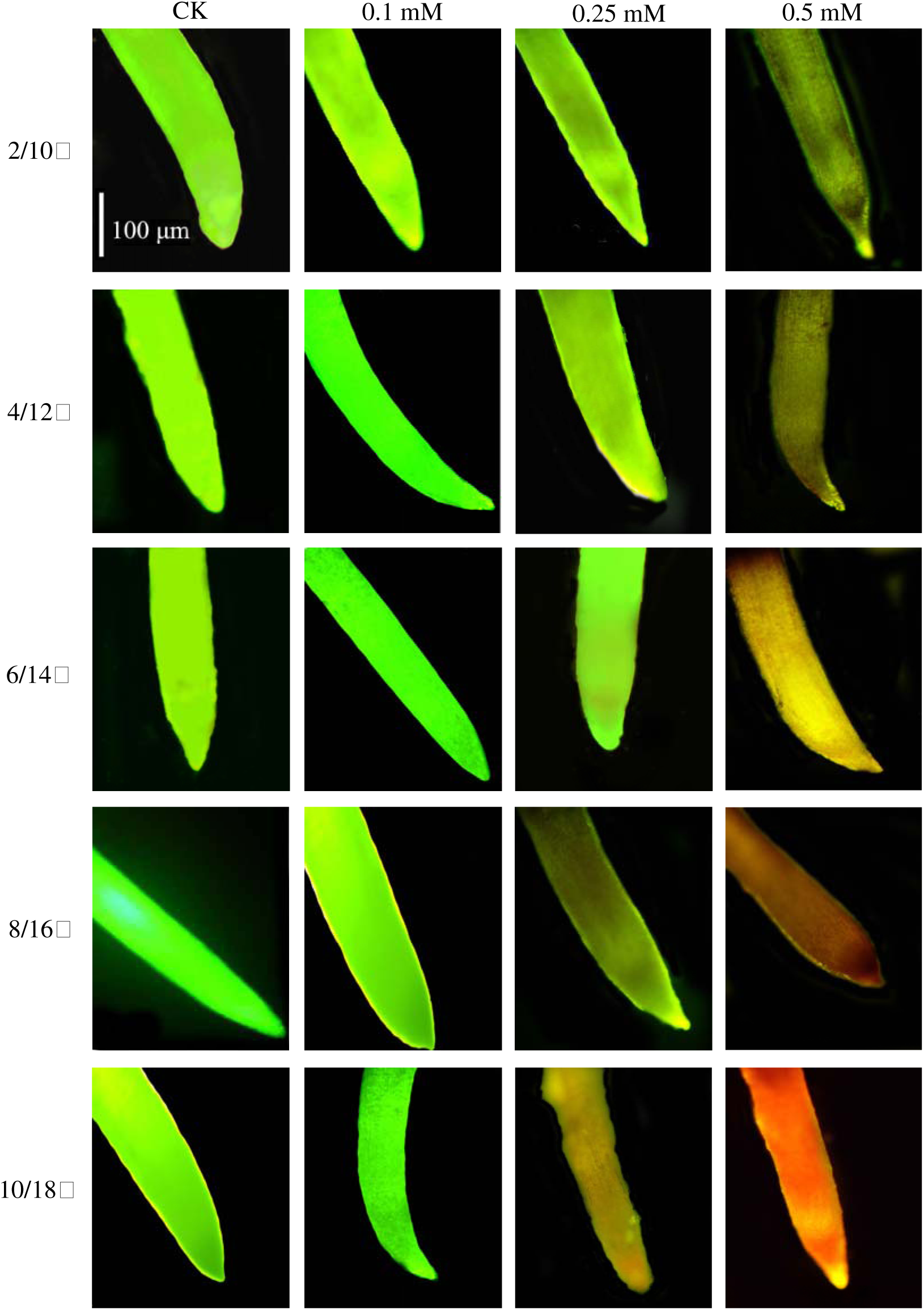
The effect of DHAP on root tips viability of *P. schrenkiana* seedlingsin different temperature ranges tested by FDA-PI staining

### Dynamic variability of auto-allelopathic effect on the antioxidant activity of enzymes

The vitality of *P. schrenkiana* seedlings is closely related to its physiological property, and thus the synergistic effects of DHAP concentration, incubation temperature and time on the activities of antioxidant enzymes in *P. schrenkiana* seedlings have been investigated. As displayed in Fig. 6, the original activities of four antioxidant enzymes SOD, POD, CAT and GR are about 45 U·g^−1^, 39 U·g^−1^, 11 U·g^−1^ and 27 μM NADPH·g^−1^ respectively (*p*<0.05), while these activities varied significantly with DHAP concentration, incubation temperature and time.

**Fig. 6.**
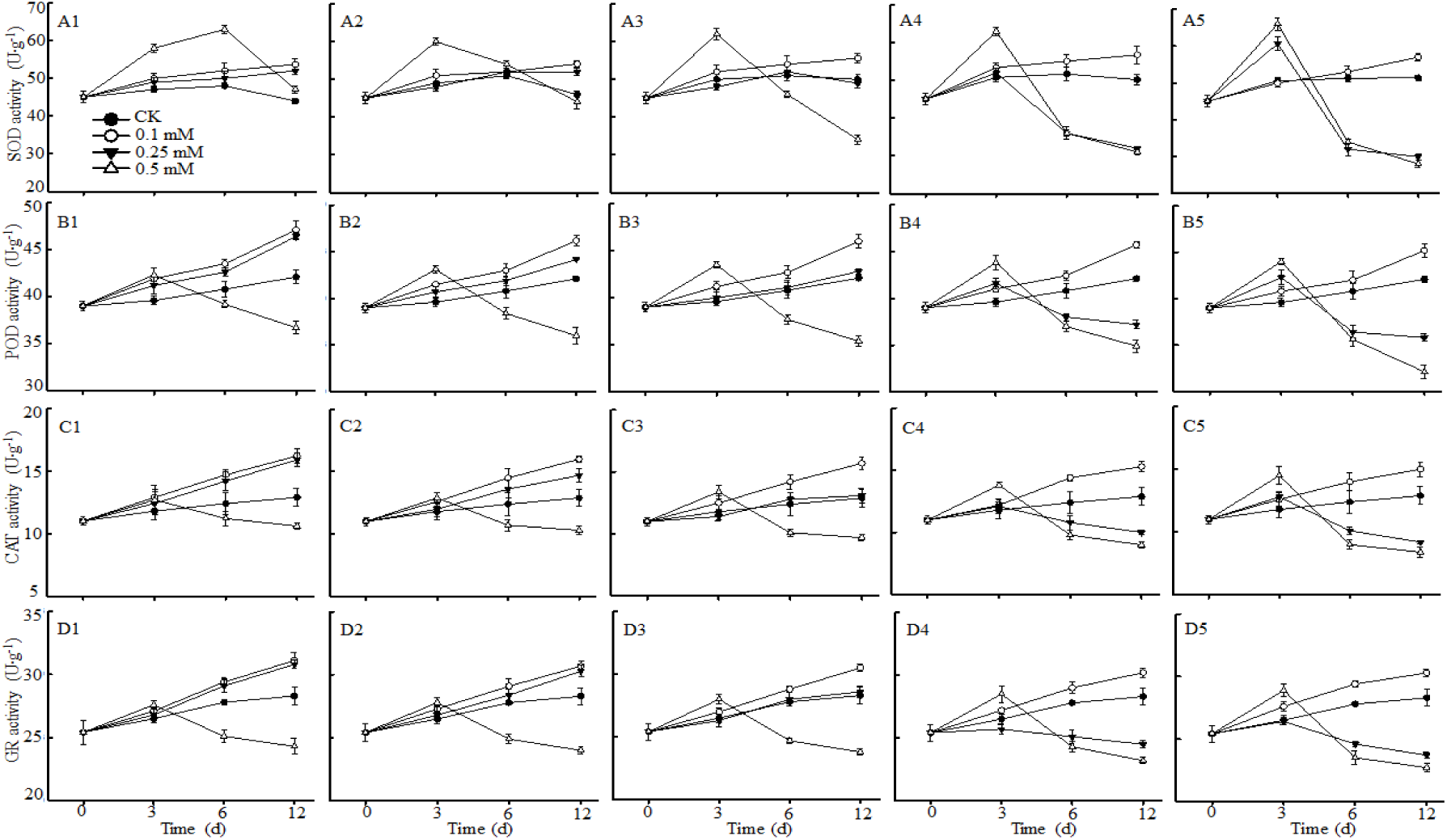
The change of antioxidant enzymes activities in *P. schrenkiana* seedlings treated by DHAP at different temperature. A1-A5: DHAP on SOD activity at 2/10°C, 4/12°C, 6/14°C, 8/16°C, and 10/18°C; B1-B5: DHAP on POD activity at 2/10°C, 4/12°C, 6/14°C, 8/16°C, and 10/18°C; C1-C5:DHAP on CAT activity at 2/10°C, 4/12°C, 6/14°C, 8/16°C, and 10/18°C; D1-D5: DHAP on GR activity at 2/10°C, 4/12°C, 6/14°C, 8/16°C, and 10/18°C

For SOD without DHAP treatment as showed in A1-A5 of Fig. 6, the intrinsic activity increased with temperature during 0 to 6 days, and then slightly descended as time extended to 12 days at temperature below 8/16 °C or approached to a stable level at 10/18 °C (*p*<0.05). For SOD with the treatment of DHAP at different concentrations and temperatures, the activity corresponding to 0.1 mM DHAP went over the intrinsic activity at any temperature and gently increased with time (*p*<0.05); the activity corresponding to 0.25 mM DHAP was higher than the intrinsic activities at 2/10 and 4/12 °C and increased with time, and was similar to the intrinsic activity at 6/14 °C and any time, while those at 8/16 and 10/18 °C went up in first 3 days and then quickly dropped down to far below the intrinsic activity after 6 days (*p*<0.05); also the activity corresponding to 0.5 mM went up significantly over the intrinsic activity at 2/10 °C during first 6 days and then dropped down to slightly above the intrinsic activity in 12 days, the activity at 4/12 °C went up to a peak during the first 3 days and then fell down to close to the intrinsic activity in 12 days, while those at temperatures over 6/14 °C ascended quickly to the maximums and then descended rapidly to far below the intrinsic activity after 6 days (*p*<0.05)..

For POD, CAT and GR without DHAP treatment, their activities at any temperature generally increased over time, as displayed by the diagrams B1-B5, C1-C5 and D1-D5 in Fig. 6 respectively. For the three enzymes with DHAP treatment, the trends and patterns of individual activity change with DHAP concentration, incubation temperature and time are basically similar to those of SOD, regardless of the difference in change extent and displacement of some turning points. For instance, their activities corresponding to 0.5 mM went up over the intrinsic activity at 2/10 °C in 3 days instead of 6 days referring to SOD and then dropped down to far below rather than slightly above the intrinsic activity referring to SOD in 12 days (*p*<0.05), as showed in A1-D1 of Fig.6.

In conclusion, the allelopathic effect of DHAP on the activities of four antioxidant enzymes in *P. schrenkiana* seedlings similarly displayed the duality relying on DHAP concentration and temperature, while time might be able to adjust the degree or even direction of such dual effects. Compared with the growth of *P. schrenkiana* seeds, seedlings and roots, however, the response of these antioxidant enzymes to the duality of DHAP allelopathic effect appeared to show some time delay. In other words, whether the promotional effect at lower concentration and temperature or the inhibitory effect at higher concentration and temperature generally required enough time to significantly change the activities of these antioxidant enzymes.

### Dynamic variability of auto-allelopathic effect on the endogenous metabolism of hormones

There are further interests to investigate the variability of DHAP allelopathy to the endogenous metabolism in *P. schrenkiana*, and hence the joint effects of DHAP concentration, incubation temperature and time on the contents of four hormones including ZT, GA_3_, IAA and ABA in *P. schrenkiana* seedling have been examined. As demonstrated in Fig.7, the contents of the four hormones in *P. schrenkiana* seedling also varied significantly with DHAP concentration, temperature and time. By comparison, the original contents of the hormones IAA, ZT, GA_3_ and ABA are about 18, 2.8, 12 and 0.01ug·g^−1^ respectively (*p*<0.05), and the change patterns of IAA, ZT and GA_3_ content show some similarities but are obviously different from that of ABA content.

**Fig. 7.**
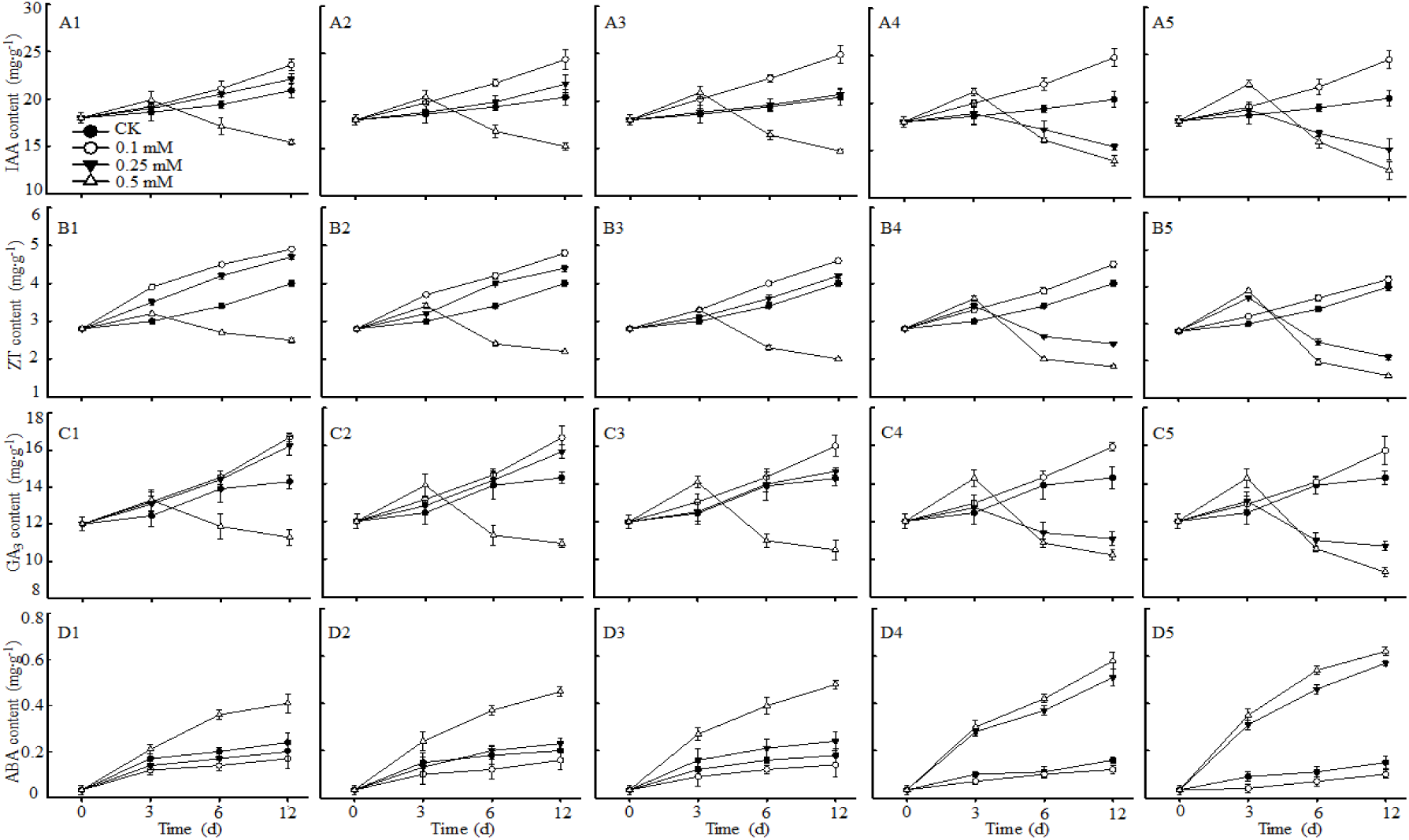
The change of endogenous plant hormones in *P. schrenkiana* seedlings treated by DHAP at different temperature. A1-A5: DHAP on IAA content at 2/10□, 4/12□, 6/14□, 8/16□, and 10/18□; B1-B5: DHAP on ZT content at 2/10□, 4/12□, 6/14□, 8/16□, and 10/18□; C1-C5: DHAP on GA3 content at 2/10□, 4/12□, 6/14□, 8/16□, and 10/18□; D1-D5:DHAP on ABA content at 2/10□, 4/12□, 6/14□, 8/16□, and 10/18□

For hormone IAA in the absence of DHAP, its intrinsic content slowly increased with time at any temperature, as showed by A1-A5 in Fig.7. In the existence of DHAP at different concentrations, the content of IAA responding to 0.1 mM DHAP rapidly increased with time at any temperature and went to much higher than the intrinsic content at 10/18 °C and 12 days (*p*<0.05); the content responding to 0.25 mM DHAP was slightly higher than the intrinsic content at 2/10 and 4/12 °C and gently increased with time, that at 6/14 °C was almost the same as the intrinsic value at any time, while those at 8/16 and 10/18 °C slowly climbed in first 3 days and then quickly slid down to far below the intrinsic values after 6 days (*p*<0.05); also the content responding to 0.5 mM DHAP went up to a peak above that responding to 0.25 mM in 3 days at any temperature and then rapidly fell down to far below the intrinsic content after 6 days (*p*<0.05).

For hormone ZT in the absence of DHAP, its intrinsic content gradually increased with time at any temperature, as showed by B1-B5 in Fig.7. In the existence of DHAP at different concentrations, the content of ZT responding to 0.1 mM DHAP quickly increased with time and went up over the intrinsic content at any time and temperature (*p*<0.05); the content responding to 0.25 mM DHAP was higher than the intrinsic content but lower than that responding to 0.1 mM DHAP at 2/10 or 4/12 °C and increased with time, and slightly higher than the intrinsic content at 6/14 °C and any time, while those at 8/16 and 10/18 °C slowly climbed in first 3 days and then quickly slid down to far below the intrinsic values after 6 days (*p*<0.05); also the content responding to 0.5 mM DHAP went up to a peak higher than that responding to 0.25 mM in 3 days at temperature above 6/14 °C and then rapidly fell down to far below the intrinsic content after 6 days (*p*<0.05).

For hormone GA_3_ in the absence or existence of DHAP as showed by C1-C5 in Fig.7, the patterns and trends of its content change with DHAP concentration, incubation temperature and time are almost the same as those of IAA, and also roughly similar to those of ZT.

For hormone ABA in absence or existence of DHAP as showed by D1-D5 in Fig.7, however, the pattern and trend of its content changing with DHAP concentration, incubation temperature and time is obviously different from those of the above three hormones. As seen, its intrinsic content and those responding to three concentrations of DHAP generally increased with temperature and time. In detail, the content responding to 0.1 mM DHAP is slightly lower than the intrinsic content at any temperature and time (*p*<0.05); that responding to 0.5 mM DHAP is much higher than the intrinsic content at any temperature and time, while that responding to 0.25 mM at any time is slightly lower than the intrinsic content at 2/10 °C, close to the intrinsic content at 4/12 °C, slightly higher than the intrinsic content at 6/14 °C, and significantly higher than the intrinsic content but slightly lower than the content responding to 0.5 mM DHAP at 8/16 and 10/18 °C (*p*<0.05). Contrary to the cases of IAA, ZT and GA_3_, in particular, low concentration of DHAP (0.1 mM) can slightly reduce the ABA content at any temperature and time, while high concentration of DHAP (0.5 mM) can significantly enhance the ABA content without a sudden and sharp drop at high temperature and any time (*p*<0.05).

In brief, the allelopathic effect of DHAP on the contents of four hormones in *P. schrenkiana* seedlings also demonstrated the duality depending on DHAP concentration and temperature, while time might magnify the extent of influence in most occasions. In general, low concentration of DHAP could gently enhance the contents of IAA, ZT and GA_3_ but slightly reduce that of ABA at any temperature and time, while high concentration of DHAP could significantly reduce the contents of IAA, ZT and GA_3_ but enhance that of ABA at high temperature and long time.

## Discussion

Global warming in conjunction with other environmental stressors and various biotic or abiotic interferences is increasingly jeopardizing the ecosystem of boreal forests (Gauthier *et al*., 2015). The existence of allelopathy has been gradually recognized by plant ecologists and chemists, but as a potential and invisible force to drive the succession of boreal forest (Inderjit *et al*., 2011), its vital role in regeneration of various plants such as *P. schrenkiana* have not attracted widespread attention, and in particular, its dynamic mechanism and ecological significance have not been fully understood yet. The main reason for that is due to the lack of innovative thinking and scientific methods. So far, most filed investigations have mainly focused on phenotypic and morphologic changes of plants (Zuo *et al*., 2007; Farooq *et al*., 2014), but hardly enter into the allelochemistry of plant secondary metabolites due to the complexity and invisibility. On the other hand, few studies in lab could clearly explain allelopathy in natural settings due to the difficulties in adopting representative samples, simulating geographical climate and ecological environment in the field, indentifying key allelochemicals and quantitatively evaluating allelopathic effects. In this study, we established a scientific methodology to simultaneously probe morphologic change, biochemistry of plant primary physiologic processes and allelochemistry of plant secondary metabolites under the simulated field conditions, accurately identify and quantitatively evaluate the allelopathic effects on plant growth, and insightfully reveal the synergistic effects of allelopathy, climate and other factors on plant behavioral ecology, Such approach may overcome the weakness of empirical, phenomenological and impractical studies in this area. As a result, this paper provides a replicable and successful example to comprehensively explain temperature-dependence of DHAP allelopathy duality and its influence on the growth of *P. schrenkiana* seeds and seedlings.

From a scientific point of view, the terminology “duality” generally refers to a basic feature of the interactions among various processes such as forests evolution. Undoubtedly, the accumulation of allelopathic substances in soil can potentially play an important role in plant adaption to environment and forest evolution. A lower cumulative amount of allelochemicals such as DHAP of 0.1 mM in the soil could promote the germination of seeds, accelerate the growth rate of seedlings, and consequently enhance the survival probability and competition level of *P. schrenkiana* population in the early stage of regeneration. By contraries, however, a higher cumulative amount of allelochemicals such as DHAP of 0.5 mM in the soil could inhibit the germination of seeds, decelerate the growth rate of seedlings, and thus reduce the survival probability and competitive ability of *P. schrenkiana* population, just appearing a “drive away” effect on the regenerated population. At the same time, the periodic fire combustion may play the role of a surface cleaner to reduce the accumulated amount of allelochemicals in topsoil and maintain the stability and balance of ecosystem (Wang *et al*., 2006). Originated from scientific thinking here, we have fully explored and quantitatively evaluated the dual effects of key alleochemical DHAP in the exudates of *P. schrenkiana* needles on the growth of its seed and seedling as well as the endogenous metabolism and antioxidant enzyme activities. In the first, an inflection concentration of DHAP was identified at any given temperature, and then it was demonstrated that depending on its concentration below or over this inflection point, DHAP might give a promotional or inhibitory effect on the growth of *P. schrenkiana* seeds and seedlings. In the second, the temperature-dependence of such DHAP allelopathic effects was verified to indicate that increasing temperature would inevitably shift the inflection concentration of DHAP from a higher level to a lower level. Consequently, these findings on the duality of DHAP allelopathy not only give a specific annotation to the universality of natural development process from quantitative change to qualitative change, but also provide a scientific guidance for the prospective researches and practices in the evolution of forest ecosystem in the context of global warming.

In the typical case given here, a scientific explanation to the temperature dependence of DHAP allelopathy duality and its influence on the growth of *P. schrenkiana* seeds and seedlings expands the understandings of allelopathic effects on forest evolution during the process of global climate change, which will help us penetrate through the apparent interaction between *P. schrenkiana* regeneration and climate change to find the common mechanism of morphological, biochemical, and ecological changes in forest evolution under global warming. Finally, this paper demonstrates our aspirations and efforts to scientifically explore the real problems of boreal forest degradation by integrating field investigation with experimental analysis and simulation. Beyond all doubt, with the continuous improvement of research methods and techniques, the role of allelopathy in plant-mediated interference to natural ecosystem is expected to become clearer in the coming years.

## Acknowledgments

The authors are grateful to the Natural Science Foundation of China (NSFC, Project No. 31670631), and Department of Science and Technology of Ningbo (DSTNB, Project No. 2017C110004, 2017C10017, 2017C10070) for the financial support of the work. Conceived and designed the experiments: Qiang Wang and Cun-de Pan. Performed the experiments and analyzed the data: Xiao Ruan, Li Yang, Min-fen Yu. Contributed reagents/materials/analysis tools: Zhao-hui Li, Cun-de Pan, and De-an Jiang. Contributed to the writing of the manuscript: Qiang Wang and Ying-xian Zhao.

## Abbreviations

DHAP: 3, 4-dihydroxyacetophenone
FDA: fluorescein diacetate
PI: propidium iodide
ZT: Zeatin
GA_3_: Gibberellin
IAA: Indoleacetic acid
ABA: Abscisic acid
SOD: Speroxide dismutase
POD: Peroxidase
CAT: Catalase
GR: Glutathione reductase

